# The Scramble Conversion Tool

**DOI:** 10.1101/003640

**Authors:** James K Bonfield

## Abstract

**Motivation:** The reference CRAM file format implementation is in Java. We present “Scramble”: a new C implementation of SAM, BAM and CRAM file I/O.

**Results:** The C API for CRAM is 1.5–1.7x slower than BAM at decoding, but 1.8–2.6x faster at encoding. We see file size savings of 40–50%.

**Availability:** Source code is available from http://sourceforge.net/projects/staden/files/io_lib/

## 1 INTRODUCTION

Storage capacity has been the primary driver behind the development of the CRAM format (Cochrane *et al.*, 2013). The European Bioinformatics Institute observed that when storing a sequence alignment to a known reference, bases that agree with the reference could be omitted and hence file size reduced (Fritz *et al.*, 2011). The CRAM format is a practical implementation of this idea and is a viable alternative to the earlier BAM format (Li *et al.*, 2009). CRAM is now the preferred submission format for the European Nucleotide Archive.

The initial CRAM prototype was in Python, quickly followed by a Picard (Wysoker *et al.*, 2009) compatible Java implementation (Zalunin *et al.*, 2011). We identified a need for a C implementation, which was implemented as part of the Staden Package’s (Staden *et al.*, 1999) “io lib” library.

The primary conversion tool is named Scramble. It can read and write SAM, BAM and CRAM formats using a unified API.

## 2 METHODS

We will not cover the CRAM file format here except to note that CRAM internally separates data by type before compressing with Zlib (Deutsch and Gailly, 1996). Thus we have regular blocks of quality values, blocks of sequence names and blocks of auxiliary tags, each of which may be compressed using different Zlib parameters. A key efficiency observation was that using the run-length-encoding strategy (“Z_RLE”) was considerably faster than the default strategy while also often offering slightly higher compression ratios for quality values. Note that this trick is not possible within the BAM format as all data types are interleaved within the same Zlib blocks.

Our implementation periodically samples both Z_RLE and the default strategy on quality blocks to determine the optimal method. This ensures rapid speed without loss in compression ratio.

Multi-threading is implemented using a thread pool, shared by both encoding and decoding tasks. This contrasts well when compared with Samtools which can only parallelize file encoding. It also permits the most efficient use of threads when converting between differing file formats, automatically balancing the encoder and decoder work loads. Note that our SAM encoding and decoding is single threaded.

## 3 RESULTS AND DISCUSSION

We tested our implementation against the reference Java cramtools implementation as well as existing BAM implementations in C (Samtools) and Java (Picard). The test data used was a 4x coverage of a *Homo Sapiens* sample (ERR317482) aligned by BWA, with a further test set (ERR251692) from the 1000 Genomes project listed in the Supplementary Information.

A break-down of the file size by item type within the Scramble CRAM output can be seen in Table 1. The impact of lossy compression on quality values was also tested by applying Illumina’s quantizing system that portions the 40 distinct values into 8 new bins (Illumina, 2012). This reduces the file size by 38%, however even in the reduced file the quality values still account for the bulk of the storage costs.

**Table 1.** CRAM breakdown by file percentage

| Data type | File %age (40 Q. bins) | File %age (8 Q. bins) |
| --- | --- | --- |
| CORE block | 6.5 | 10.6 |
| Substituted/insert bases | 0.0 | 0.0 |
| Soft-clip bases | 0.2 | 0.3 |
| Quality values | 79.6 | 66.9 |
| Sequence identifiers | 8.2 | 13.3 |
| Template position/size | 0.2 | 0.3 |
| Auxiliary tags | 4.9 | 8.1 |

Total file sizes: 3.51Gb for 40 bins, 2.16Gb for 8 bins.

Table 2 shows the time taken to read and write formats from the various tools along with their resultant file sizes. For encoding it is clear that the C implementation of CRAM is considerably faster than the Java implementation and also beats Picard/Samtools BAM speed despite the use of the Intel tuned Deflate implementation by Picard. This is almost entirely down to the use of Z_RLE for encoding quality values. Decoding of CRAM is not as fast as C BAM, but it is comparable to the widely used Picard’s BAM decoder. We also observe that the CRAM files produced by Scramble are around 7% smaller than those produced by Cramtools.jar.

**Table 2.** 9827 2#49.bam (ERR317482)

| Tool | Format | 40 quality bins |  |  | 8 quality bins |  |  |
| --- | --- | --- | --- | --- | --- | --- | --- |
|  |  | Read(s) | Write(s) | Size(Gb) | Read(s) | Write(s) | Size(Gb) |
| Scramble | BAM | **76.2** | 680.1 | 6.50 | **63.1** | 839.3 | 4.80 |
| Scramble | CRAM | 117.1 | **325.2** | **3.51** | 115.2 | **302.7** | **2.16** |
| Cramtools | CRAM | 212.1 | 1385.1 | 3.78 | 195.7 | 1365.1 | 2.33 |
| Samtools | BAM | 78.1 | 672.7 | 6.50 | 67.0 | 831.6 | 4.80 |
| Picard | BAM | 123.8 | 532.7 | 6.52 | 110.6 | 468.6 | 4.90 |

User + System CPU times in seconds for encoding and decoding along with the produced file size.

Scramble has full multi-threading support for both reading and writing of BAM and CRAM file formats. It scales nearly linearly up to 16 cores, but with some performance inefficiencies becoming visible in CRAM with high core counts. The results can be seen in Figure 1.

**Fig. 1.**
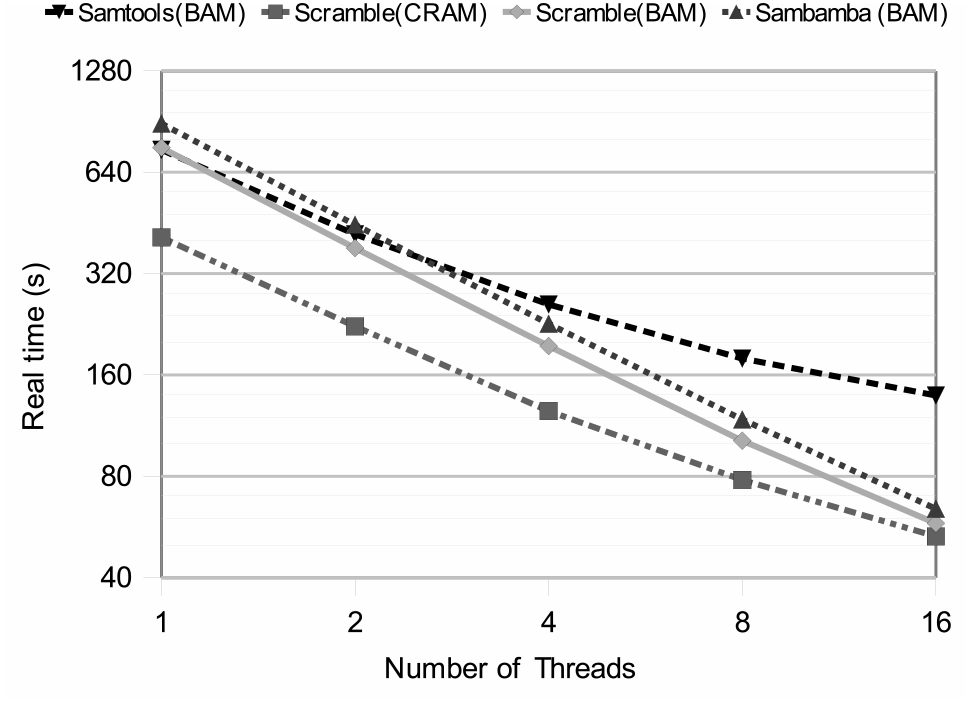
Real time taken to convert from BAM to BAM/CRAM format using Scramble and Samtools (BAM to BAM only).

## 4 CONCLUSION

We have demonstrated that the C implementation of CRAM performs well, beating Samtools, Picard and Cramtools for encoding speed. Decoding speed is not as efficient as Samtools, but is still comparable to Picard and nearly twice as fast as the Java CRAM implementation.

CRAM is not capable of achieving the top compression ratios, using 3.96 bits/base with 40 quality bins and 2.05 bits/base with 8 bins compared against only 3.16 and 1.52 for fqz comp (Bonfield and Mahoney, 2013), and 41 bits per read name in CRAM versus 23 bits in fqz_comp. This demonstrates room for improvement in future CRAM versions, possibly achieved by implementing arithmetic coding instead of relying on Zlib.

Scramble is not a drop-in replacement for the Samtools API however a port of the CRAM components of Scramble has been made to the HTSlib library and is available within a test release of Samtools, available from *https://github.com/samtools/htslib*.

## ACKNOWLEDGEMENT

We would like to acknowledge Vadim Zalunin for his assistance and collaboration with re-implementing the CRAM specification.

*Funding*: This work was funded by the Wellcome Trust [098051].

